# Assembly and translocation of a CRISPR-Cas primed acquisition complex

**DOI:** 10.1101/208058

**Authors:** Maxwell W. Brown, Kaylee E. Dillard, Yibei Xiao, Adam Dolan, Erik Hernandez, Samuel Dahlhauser, Yoori Kim, Logan R. Myler, Eric Anslyn, Ailong Ke, Ilya J. Finkelstein

## Abstract

Bacteria and archaea destroy foreign nucleic acids by mounting an RNA-based CRISPR-Cas adaptive immune response^1–3^. In type I CRISPR-Cas systems, the most frequently found type of CRISPR in bacteria and archaea^3,4^, foreign DNAs that trigger efficient immunity can also provoke primed acquisition of protospacers into the CRISPR locus^5–12^. Both interference and primed acquisition require Cascade (CRISPR-associated complex for antiviral defense) and the Cas3 helicase/nuclease. Primed acquisition also requires the Cas1-Cas2 integrase; however, the biophysical mechanisms of how interference and primed acquisition are coordinated have remained elusive. Here, we present single-molecule characterization of the type I-E *Thermobifida fusca (Tfu)* primed acquisition complex (PAC). *Tfu*Cascade rapidly samples non-specific DNA for its target via facilitated one-dimensional (1D) diffusion. An evolutionary-conserved positive patch on the Cse1 subunit increases the target recognition efficiency by promoting this 1D diffusion. Cas3 loads at target-bound Cascade and the Cascade/Cas3 complex initiates processive translocation via a looped DNA intermediate. Moving Cascade/Cas3 complexes stall and release the DNA loop at protein roadblocks. Cas1-Cas2 samples DNA transiently via 3D collisions, but stably associates with target-bound Cascade. Cas1-Cas2 also remains associated with translocating Cascade/Cas3, forming the PAC. By directly imaging all key subcomplexes involved in target recognition, interference, and primed acquisition, this work provides a molecular basis for the coordinated steps in CRISPR-based adaptive immunity.

## Introduction

CRISPR-Cas adaptive immunity consists of three main processes: interference, primed acquisition, and naïve acquisition^13–16^. Interference provides immunity by targeting and destroying foreign nucleic acids whose sequences are recorded in the CRISPR array. During acquisition, the CRISPR system adapts to new threats by incorporating segments of foreign genetic elements into the CRISPR array, where they are then transcribed and used to confer immunity against the invading nucleic acid^17–20^. In Type I-E CRISPR-Cas systems, the Cascade surveillance complex, consisting of 11 subunits of five Cas proteins (Cse1_1_, Cse2_2_, Cas7_6_, Cas5_1_, and Cas6e_1_) and a 61-nucleotide (nt) CRISPR RNA (crRNA), initiates both interference and primed acquisition^5–12^. Cascade surveys the cell for foreign DNA that is complementary to its crRNA^17^. An RNA-DNA loop (R-loop) between the crRNA and the duplex target DNA conformationally locks Cascade onto the foreign genetic element^21–29^. Next, target-bound Cascade loads Cas3 nuclease/helicase, which unwinds and degrades the foreign DNA into a single-stranded DNA (ssDNA) product^21,30–33^.

Primed and naïve acquisition both require the Cas1-Cas2 integrase. Cas1-Cas2 inserts new protospacers into the CRISPR locus in the host’s genome via a cut-and-paste transposase mechanism^34–36^. Naïve acquisition can integrate foreign nucleic acids that the cell has not encountered previously and requires host nucleases to produce substrates for Cas1-Cas2^37^. In contrast, primed acquisition uses Cascade/Cas3 to produce protospacers that Cas1-Cas2 then integrates into the CRISPR locus^5–9,38,11,10^. Primed acquisition thus requires a prior record of infection by a related pathogen. Because primed acquisition is substantially more efficient than naïve acquisition, this mechanism permits the cell to rapidly adapt to phages that have acquired escape mutations^5,9,11,39^. Although the genetic requirements for primed acquisition have been established previously, the biophysical mechanisms underpinning interactions between Cascade, Cas3, and Cas1-Cas2 have remained elusive^5,6,9^. To address this gap, we report the stepwise assembly and biophysical characterization of the *Thermobifida fusca (Tfu)* type I-E CRISPR-Cas interference and primed acquisition machineries. Using single-molecule fluorescence imaging of each subcomplex, we show that Cse1, a subunit of Cascade, plays a key role in target recognition by facilitating rapid scanning of foreign DNA via facilitated diffusion. After target recognition, Cascade recruits Cas3, and the Cascade/Cas3 interference complex translocates via a looped DNA intermediate. Finally, we provide direct evidence that Cascade/Cas3 interacts with Cas1-Cas2 to form a translocating complex that combines all the biochemical functions required for both interference and primed acquisition.

## Results

### Cse1 promotes target recognition via facilitated diffusion on nonspecific DNA

To understand how Cascade participates in both interference and primed acquisition, we first imaged fluorescent *Tfu*Cascade on double-tethered DNA curtains that extend the substrate in the absence of buffer flow^40,41^ (Figures 1 and S1). The DNA substrate (derived from bacteriophage λ) lacked a target DNA sequence that was complementary to the Cascade crRNA. Prior studies reported that the *S. pyogenes* Cas9 and *E. coli* Type I-E effector complexes sample protospacer adjacent motif (PAM) sites exclusively via three dimensional (3D) collisions, suggesting that this is a universal feature of diverse CRISPR systems^42,43^. Unexpectedly, 90% (N=258 out of 288) of *Tfu*Cascade molecules initially bound non-specific DNA and scanned the substrate via facilitated one-dimensional (1D) diffusion (Figure 1D-F). Facilitated diffusion can accelerate the target search dynamics, as has been observed for both DNA and RNA-binding proteins^44,45^. Proteins that scan DNA via 1D diffusion can either slide along the helical pitch of the DNA backbone, or can transiently dissociate and associate with the DNA via a series of microscopic hops. Hopping allows proteins to efficiently search larger segments of the genome, while frequently randomizing the spatial register between the protein and the DNA backbone (see below)^44,46^. Hopping can be observed indirectly by measuring the change in the diffusion coefficients at higher ionic strengths, which increases electrostatic screening between the protein and DNA. This results in measurably larger 1D diffusion coefficients and can be used to estimate the number of disrupted electrostatic charges^47^. Cascade diffusion coefficients increased with higher ionic strength, indicating that both sliding and hopping mechanisms contribute to target scanning (Figure 1F). Cascade lacking Cse1 did not diffuse on DNA curtains. Therefore, we conjectured that a positive patch on the *Tfu*Cse1 outer surface (Figure 1A, bottom) promotes facilitated diffusion of Cascade during foreign DNA surveillance and that increasing ionic strength screens at least one of these charges (Figure 1F). A structure-based multi-sequence alignment of divergent Cse1 variants revealed that the positive patch is highly conserved and can extend up to eight amino acids (Figure S2A)^48,49^. Notably, this positive patch is disrupted in the *E. coli (Ec)* Cse1, likely limiting the 1D scanning mode of *Ec*Cascade beyond the resolution of prior studies (Figure S2B)^42^. The *Tfu*Cse1 studied here encodes positive charges at five of these eight sites (Figure 1A). To test the importance of the Cse1 positive patch on facilitated diffusion, we purified Cascade harboring Cse1(5A), a variant with all five positive residues mutated to alanine (Figure S2B). Cse1(5A)-Cascade diffusion trajectories were 2.6-fold shorter than the wild type complex on non-specific DNA (Figure 1G; 2.7 ± 0.7 sec, N=50 molecules vs. 7.1 ± 1.8 sec, N=100), and also had a 50-fold lower binding affinity for target DNA, as determined by electrophoretic mobility shift assays (EMSAs, Figures 1H and S2). Extending the positive patch to eight positive residues, Cse1(3R), did not appreciably change the duration of the diffusion traces (8.9 ± 2.2 sec, N=100) and also did not affect the binding affinity for target DNA (Figures S2), indicating that additional charges are not necessary for efficient target recognition. To further probe the role of Cse1 in promoting Cascade diffusion, we optimized a sortase-based transpeptidation strategy to fluorescently label the Cse1 subunit alone, or in complex with Cascade (Figure S3)^50^. Fluorescent Cse1 could bind and diffuse on DNA, with the longest Cse1 binding events occurring on DNA regions with the highest PAM density (Figure S3D-F). Cse1 diffusion trajectories were shorter than those for the Cascade complex at identical ionic strength, suggesting that Cascade also contributes secondary non-specific DNA interactions (Figure S3E). A positive groove in the *Tfu*Cse2 subunit is positioned to interact with DNA in the Cascade-crRNA structure and may contribute additional stabilization during target search on non-specific DNA^27,51–53^. Taken together, this data shows that the positive channel formed on the surface of *Tfu*Cse1 is critical in promoting facilitated diffusion and efficient target recognition by *Tfu*Cascade.

**Fig. 1.**
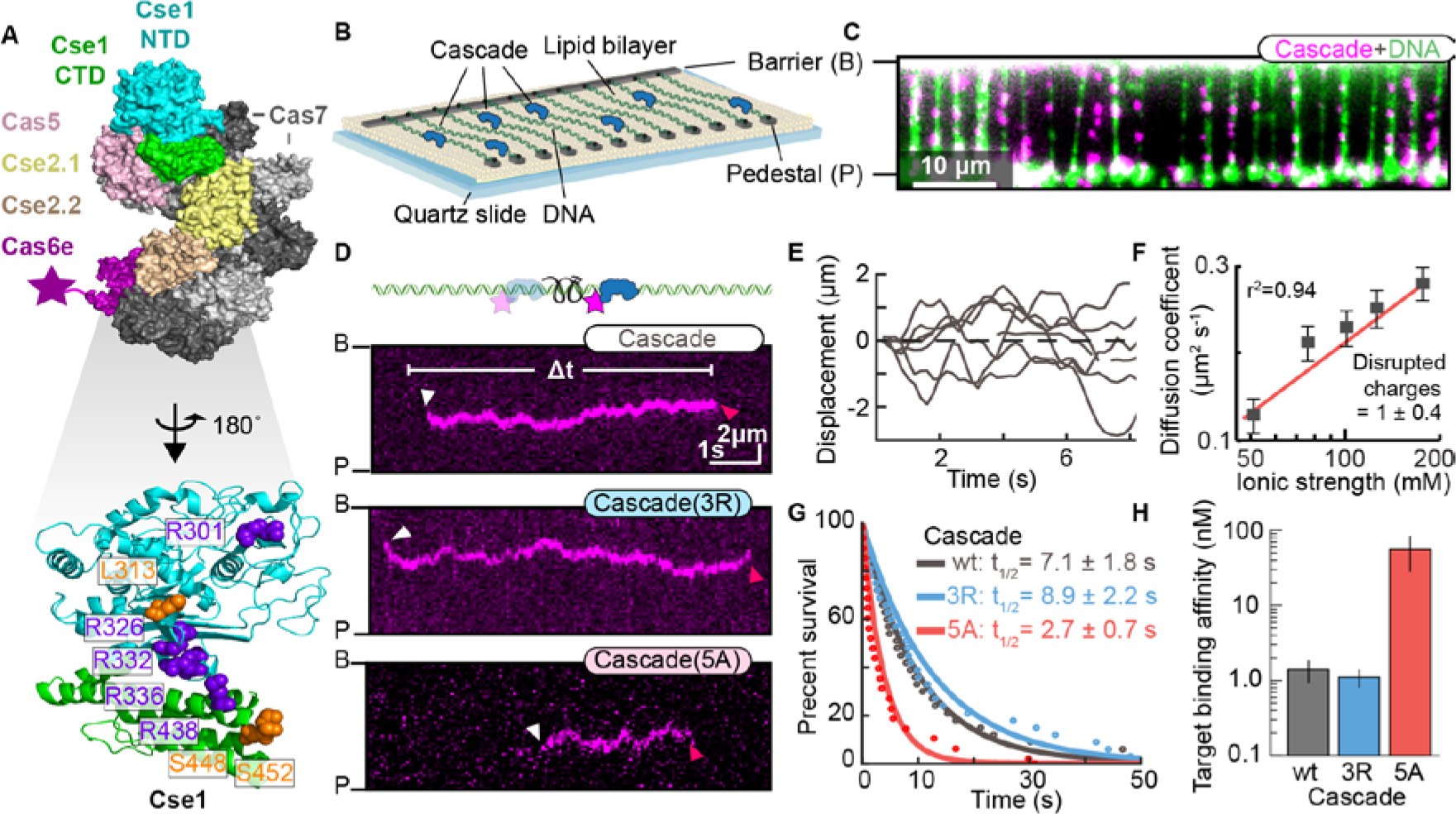
Cse1 promotes facilitated diffusion of the Cascade surveillance complex along DNA. (A) Top: structure of the *T. fusca* (*Tfu*) Cascade surveillance complex (PDB ID: 5U0A). An epitope on the C-terminus of Cas6e was used for fluorescent labeling (star). Bottom: an evolutionarily conserved positive patch that is conserved (purple) and neutral (orange) in *Tfu*Cse1. (B) Illustration of double-tethered DNA curtains. A lipid bilayer is deposited on a quartz slide with a microfabricated chrome barrier (B) and pedestals (P). Phage λ DNA is ligated with biotin and digoxigenin (dig)-terminated oligonucleotides and tethered to the lipid bilayer via a biotin-streptavidin linkage. The second DNA end is immobilized on pedestals coated with anti-digoxigenin antibodies. (C) Fluorescent image of double-tethered DNA curtains. DNA is stained with a fluorescent intercalating dye (YOYO-1, green). Cascade (magenta) binds non-specifically along the DNA substrate. (D) Illustration (top) and kymographs (bottom) of the indicated Cascade variants scanning DNA for targets via facilitated diffusion. White and red arrows mark DNA binding and release, respectively. (E) Single-particle traces showing six representative Cascade molecules diffusing on DNA. (F) Mean Cascade diffusion coefficients as a function of the ionic strength. N > 45 molecules for all conditions. Error bars: S.E.M. The linear fit (red line) estimates 0.93±0.43 (Avg ± 95% C.I.) Coulombic interactions are disrupted at increasing ionic strength. (G) DNA-binding lifetimes of each Cascade variant. The data was fit to a single exponential decay (solid lines). Half-lives ± 95% C.I. is calculated from the fit. (H) Cascade target binding affinities, as measured via electrophoretic mobility shift assays. Mean and S.D. are calculated from at least three replicates.

### Cascade samples potential targets via two transient intermediates

Next, we determined how diffusing Cascade molecules recognize full and partially complementary DNA targets (Figure 2). Incubating Cascade with a target-containing DNA substrate prior to imaging resulted in complexes that remain bound at the target site for > 1,900 seconds, indicating full R-loop propagation and R-loop locking (Figures 2B & S4)^22,28,42,54^. Next, we imaged the target search and recognition reaction by adding fluorescent Cascade to pre-assembled DNA curtains. Surprisingly, diffusing complexes frequently paused and released the target site without forming a stable R-loop (Figure 2C). Approximately 80% of Cascade-target encounters (N=313 encounters) resulted in pausing and release events (defined to be >800 ms for high-confidence pauses; see methods) (Figure 2D). Diffusing Cascade only pauses at full or partial targets; we did not observe pausing on PAM-rich, but otherwise non-specific DNA (see next paragraph). Cascade first recognizes the PAM via the Cse1 subunit, followed by directional extension of the R-loop away from the PAM and along the crRNA^55,54,28,22,43^. However, diffusing Cascade can encounter the target in two polarities—with Cse1 positioned to recognize the PAM and the crRNA oriented in the correct direction for R-loop propagation, or with the crRNA in the opposite orientation relative to the target DNA. Therefore, we also determined whether Cascades that encounter the target from the PAM-proximal or distal sites impact the target recognition frequency. Remarkably, complexes that approach from the PAM-proximal side were just as likely to pause at the target site as those that approach from the PAM-distal end (Figure 2D). Moreover, after the pauses, Cascade was equally likely to depart from the target site in either PAM-proximal or distal direction (Figure S4C). This orientation-independent target recognition is consistent with microscopic hopping during facilitated diffusion. Hopping allows Cascade, and likely other site-specific DNA binding proteins, to sample potential target sites with both polarities, ensuring efficient target recognition.

**Fig. 2.**
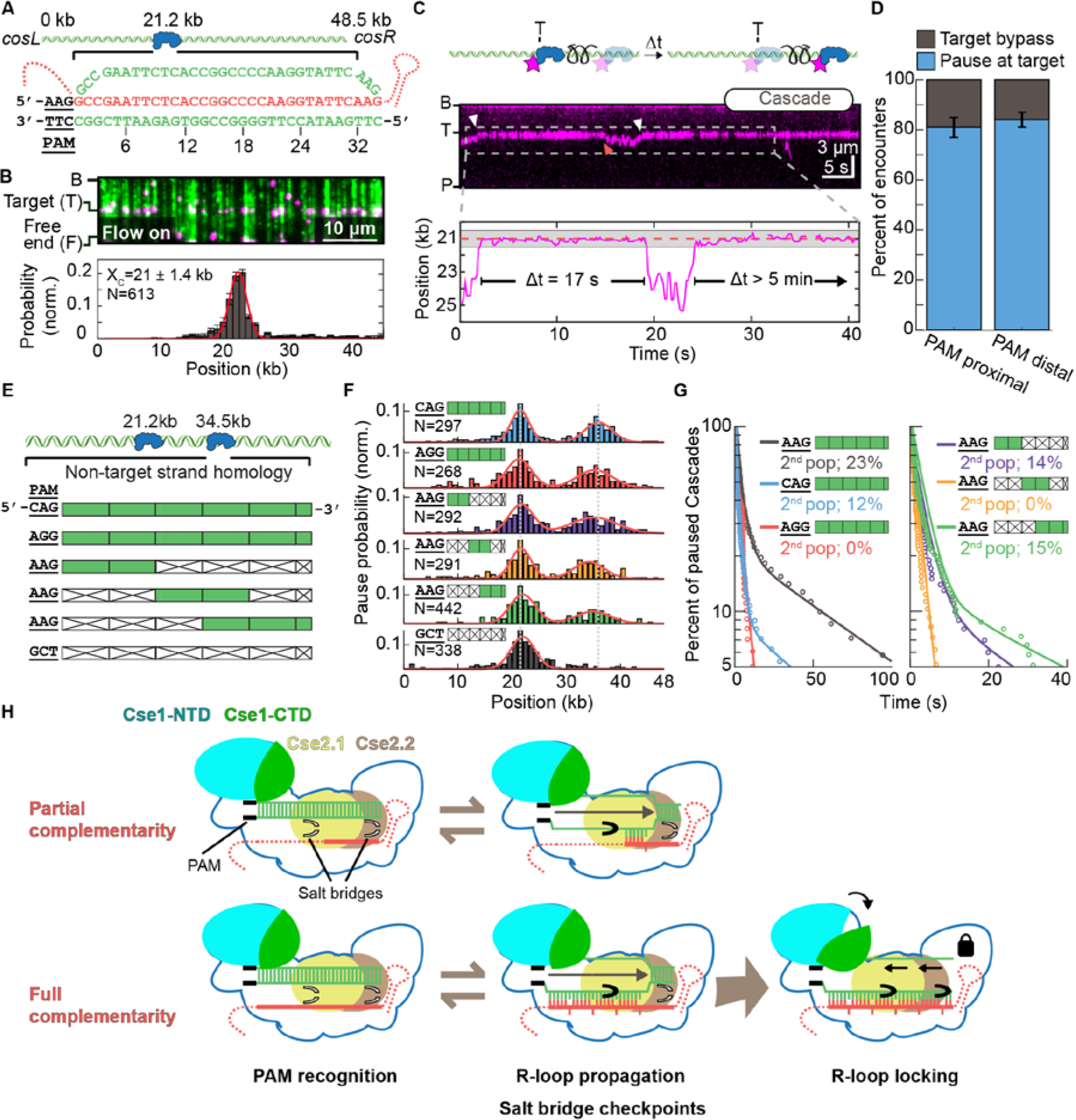
Cascade transiently samples target sequences via PAM-dependent R-loop propagation and seed-distal complementarity. (A)Illustration of a DNA substrate with a single Cascade target inserted 21.2 kb away from the *cosL* DNA end. The target DNA strand is shown base-paired to the crRNA (red). Numbers indicate flipped out R-loop bases. (B) Top: image of Cascade (magenta) bound to the target sequence on a singletethered DNA curtain (green). Bottom: histogram of Cascade binding the target site. Red line: Gaussian fit with the center and standard deviation of the fit (error bar) indicated in the figure. (C) Top: illustration and kymograph of a diffusing Cascade molecule transiently pausing at the target site. The white and red arrows indicate the beginning and end of a pause, respectively. Bottom: single-molecule tracking indicates that Cascade pauses twice at the target site (dashed line). The gray band indicates the experimental uncertainty in defining the target site. (D) Cascade pauses with equal frequency at the target regardless of whether it approaches from the PAM-proximal or PAM-distal side (N=27 Cascade molecules; 227 pauses). Error bars are generated via bootstrapping. (E) Schematic of six DNA substrates containing a second Cascade target 34.5 kb away from the *cosL* DNA end. The second targets encode either an altered PAM or segments of the target DNA that are mismatched (white boxes) or complementary (green boxes) to the crRNA. The bottom DNA substrate does not encode extensive complementarity to the crRNA and is included as a negative control. (F) Pausing probability of Cascade on the six DNA substrates described in (E). Pausing distributions are fit to two Gaussians (red) and recover both target positions (dotted grey lines). N: number of pauses. (G) Cascade pause durations on the substrates shown in (E). In all but two cases, the data required a bi-exponential fit (solid lines). The magnitude of the second population of the two exponentials is reported. N > 95 pauses for all experiments. (H) Model for target recognition by diffusing Cascade surveillance complexes. Cse1 interacts with the PAM to begin directional unwinding of the DNA duplex. Top row: the extending R-loop is partially stabilized by salt bridges in *Tfu*Cse2.1 and Cse2.2 (black closed gates)^27^, but eventually collapses, causing Cascade to leave the target. Bottom: complete R-loop extension locks Cascade onto the DNA target, triggers a conformational change in Cse1, and promotes Cas3 binding (not shown).

Cascade is proposed to engage potential target DNA sites via a series of sequential steps that include PAM recognition, melting of a DNA bubble, propagation of the R-loop past a critical 8-10 nt ‘seed’ region, and conformational locking^6,22,26–28,54,55^. To further probe this series of steps, we constructed DNA substrates that included a second target site at 34.5 kb with altered PAMs or partial sequence complementarity to the crRNA (Figures 2E and S4D). Cascade pausing at these partial target sites required both a PAM as well as a segment of target DNA complementary to the crRNA. Surprisingly, scrambling the seed region only resulted in a 50% reduction of paused Cascade molecules relative to the perfect target sequence (Figure 2F). This suggests that Cascade can transiently recognize PAM-distal target DNA independently of the seed^22^. Next, we observed how long Cascade remained associated with each of the PAM variants and partial target sequences (Figure 2G, left). Cascade pause times were best described by a bi-exponential fit with a short, *t*_*1*_=1-3 sec, and a longer, t_2_~50 sec, half-life. The PAM controlled the duration and relative amplitude of the shorter timescale (*t*_*1*_), but not the duration of *t*_*2*_. The highest DNA-binding affinity (and strongest interference) PAM (5’-AAG) resulted in the longest *t*_*1*_ pause duration *t*_*1*_=2.8 ± 0.1 sec (N=656 pauses). In contrast, intermediate interference 5’-CAG and weakest interference 5’-AGG PAMs had short *t*_*1*_ pauses (*t*_*1*_=1.5 ± 0.1 sec; N=105 and *t*_*1*_=2.4 ± 0.4 sec; N=96 pauses, respectively). Moreover, the weakest 5’-AGG PAM pause durations were best described by a single, short exponential decay without a long-lived state (*t*_*2*_). Next, we determined the pause duration for Cascade on a series of targets that had the strongest PAM (5’-AAG), but contained mismatches between the crRNA and the first, second, and third segments of the target DNA (Figure 2G, right). All DNA substrates still exhibited a short pause, *t*_*1*_=~1-2 sec. The second pause duration, *t*_*2*_, was ~2.6 fold shorter than the perfect target for substrates with PAM-proximal and distal complementarity, but was virtually non-existent when the complementarity was moved to the middle segment. These data show that complementarity in the PAM-proximal ‘seed’ region is sufficient to induce a long-lived pause on the partial target as the R-loop directionally propagates away from the PAM. Unexpectedly, PAM-distal complementarity is also sufficient for a long-lived Cascade pause. Our recent structural snapshots of a partial *Tfu*Cascade R-loop revealed that salt-bridges between Cas7s and the two Cse2 subunits seal the target strand in PAM-distal regions during R-loop propagation^27^. Taken together, the structural and single-molecule results suggest the model summarized in Figure 2H. The identity of the PAM and the first few PAM-proximal nucleotides initiate a short (1-3 sec) pause. This pause is likely necessary for Cse1 to insert an aromatic wedge into the PAM-proximal DNA duplex and melt a bubble in the target DNA^26^. R-loop propagation is reversible, even on the complementary target DNA. Extension of the R-loop past two Cse2 salt bridges further stabilize the R-loop intermediate. Finally, conformational locking of the entire Cascade complex re-orients the Cse1 N-and C-terminal lobes for Cas3 recruitment and downstream interference and primed acquisition^25^.

### Translocating Cascade/Cas3 complexes generate tension-sensitive DNA loops

Primed acquisition and interference both require the concerted activities of Cascade and the Cas3 nuclease/helicase. Therefore, we next determined the mechanism of *Tfu*Cas3 recruitment and translocation by imaging Cascade, Cas3, and the single-strand DNA (ssDNA) product. For fluorescent imaging, an ATTO647N dye was directly conjugated to the C-terminus of Cas3 via sortase-mediated transpeptidation (Figure S5). Labeling Cas3 with a small C-terminal organic fluorophore was essential because the N-terminus of Cas3 interacts with *Tfu*Cas1-Cas2 (data not shown). Fluorescent Cas3 preferentially localized to target-bound Cascade and remained stationary on the single-tethered DNA substrates with AMP-PNP, a non-hydrolysable ATP analog (Figures 3A and S5B-D). These findings are consistent with Cascade loading Cas3 onto the target DNA. In the presence of 1 mM ATP, Cas3 translocated towards the DNA tethering point, as expected for the 3’ to 5’ directionality of the Cas3 helicase domain on the non-target strand (Figure 3B & S5)^56^. Remarkably, Cascade remained associated with the translocating Cas3 in 47% of all trajectories (Figure 3B). In the remaining trajectories, Cascade and Cas3 fluorescent signals separated within a single frame (< 200 ms), suggesting a rupture between Cascade and Cas3 that was rapid and stochastic. After rupturing from Cas3, Cascade returned to its initial position at the target DNA site while Cas3 continued to translocate along the DNA substrate (Figure 3B, top). The co-translocation of the Cascade/Cas3 complex and instantaneous Cascade return to the target site is consistent with a looped DNA intermediate produced during DNA translocation. The DNA loop is produced because Cas3 translocates away from the target site while maintaining contact with target-bound Cascade, as has been proposed for the *E. coli* Type I-E system^42,57^. Individual trajectories revealed an initiation phase where Cas3 did not appear to translocate. However, we detected limited Cas3 helicase/nuclease activity during this initiation phase via production of short ssDNA segments that could be visualized by adding fluorescent single-stranded DNA binding protein (SSB-GFP) to the flowcells. (Figure S6C). Our results are consistent with a smFRET study that detected short burst of Cas3 translocation that are below our spatial resolution^57^. This initiation phase lasted for 30 ± 0.8 s (N=48), followed by processive movement along the DNA substrate (Figure 3C). Cas3 translocated along the DNA substrate with a mean processivity of 19 ± 7 kb (N=68, error denotes S.D.) at a velocity of 89 ± 25 bp s^−1^ (N=68). Guided by previous findings that Cas3 interacts with the Cse1 subunit of Cascade and observation that Cse1 and Cas3 are fused in other type I-E systems, we tested whether Cse1 is associated with translocating Cas3 after Cascade release^31,42^. Concurrent dual-color imaging of both Cse1 and Cas6e in a dual-labeled (ATTO647N) Cse1-Cascade complex revealed that Cse1 always remained associated with Cascade as Cas3 translocated awayfrom the effector complex (Figure S5E). These results provide direct evidence for retention of Cse1 in the Cascade effector complex after Cas3 loading and translocation. Physical interactions between target-bound Cascade and a moving Cas3 will produce a growing and tension-dependent DNA loop that is extruded at the Cse1-Cas3 interface^42,57^. To directly visualize these looped DNA intermediates, we used DNA substrates with one fluorescent DNA end positioned either upstream or downstream of translocating Cas3 (Figure S5F,G). Consistent with the looping model, Cas3 movement away from the free DNA end pulls Cascade and the free DNA end at identical rates in the direction of Cas3 translocation (Figure S5F). Alternatively, if the DNA tethering geometry is reversed, then Cas3 translocation will reel in the free DNA end without observable Cascade movement (Figure S5G). Retraction and stochastic release of the free DNA end corresponded with Cas3-dependent translocation and Cse1-Cas3 rupture. In the cell, one or both ends of the foreign DNA are likely to be physically constrained (i.e., to the viral capsid during infection/package or to the transcription/translation machinery during viral replication)^58^. Processive Cascade/Cas3 translocation will thus produce increasing DNA tension as the DNA loop grows. To define the role of DNA tension on Cas3 translocation, we developed a high-throughput assay to measure force-dependent Cascade/Cas3 loop rupture (Figures 3D and S5H). In this assay, one end of the DNA is immobilized on the pedestals and the second DNA end is conjugated to 1 µm streptavidin-coated paramagnetic beads. The tension on the DNA molecule can then be modulated by increasing the force on the bead. These beads increase the hydrodynamic drag experienced by DNA molecules under mild buffer flow. Increasing the buffer flow rate (hydrodynamic force) correspondingly increases the tension applied to the DNA molecule (Figure S5I). At an applied force of 0.7 pN, 53% (N=30) of translocating Cascade/Cas3 complexes moved together as a complex for the duration for the entire trajectory. Increasing the applied force resulted in substantially fewer looped Cascade/Cas3 complexes; only 11% (N=18) of translocating complexes moved together at 20 pN of applied force (Figure 3D). We conclude that Cascade/Cas3 interactions rupture as tension accumulates between the moving Cas3 and stationary Cascade.

**Fig. 3.**
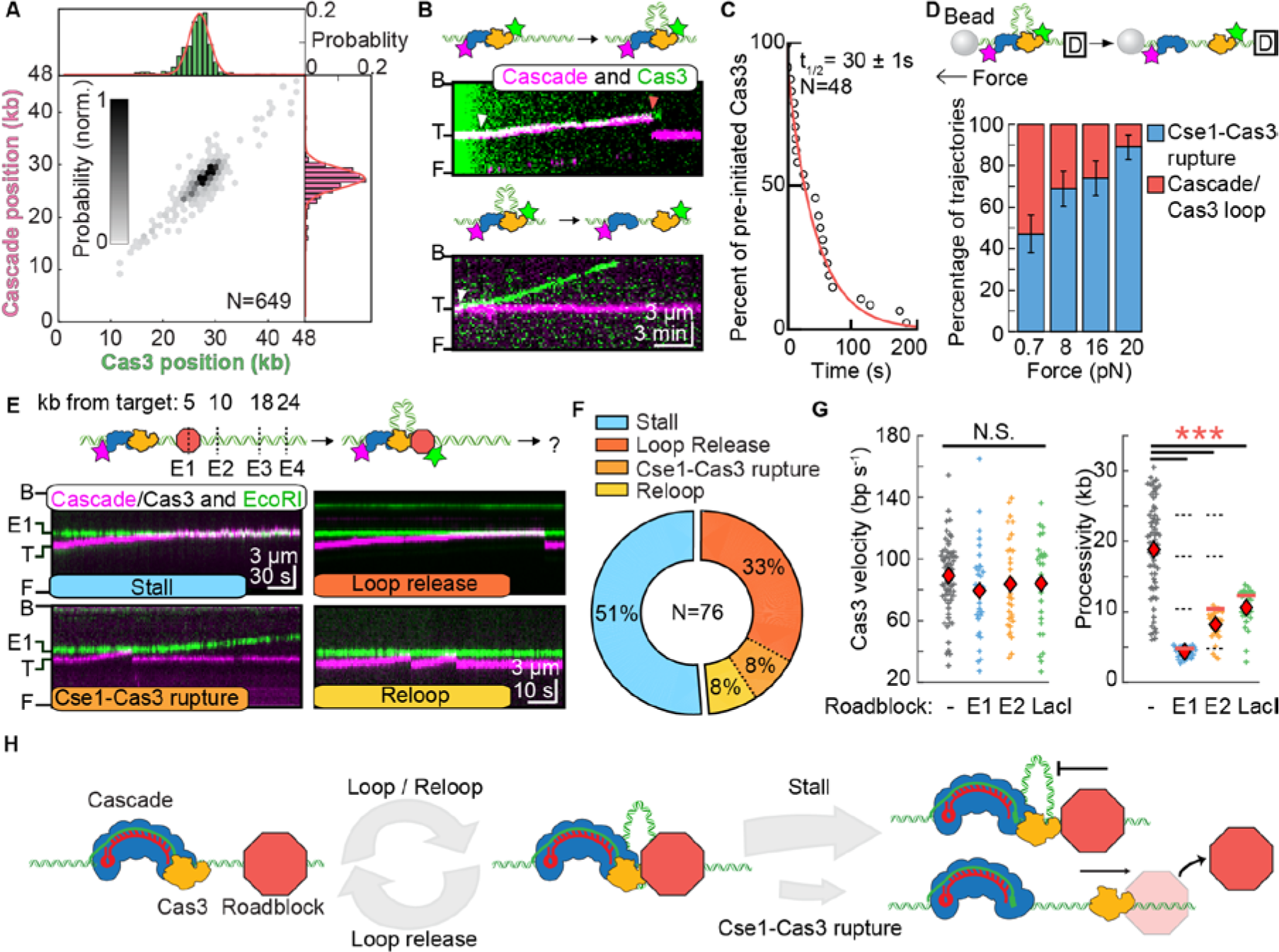
Processive translocation by the Cascade/Cas3 complex is impeded by DNA-binding proteins. (A)Histograms of Cas3 (top), Cascade (right), and their joint DNA-binding probability (center) indicate that Cascade loads Cas3 at the target site. (B) Top: illustration and kymograph of a translocating Cascade/Cas3 complex. Cascade remains associated with the target, causing a DNA loop to accumulate during Cas3 translocation. Bottom: Cas3 translocating independently of Cascade. White arrows: initiation of translocation; red arrow: Cascade/Cas3 separation. (C) Cas3 initiates translocation after a 30 ± 1 second pause (N=48). The pause data was fit to a single exponential decay (solid line) to calculate the half-life. Error indicates 95% C.I. (D) Top: illustration of force-dependent Cas3 translocation experiments. The free DNA end was conjugated to a 1 μm paramagnetic bead and hydrodynamic force was applied via buffer flow. Increasing tension on the DNA also increases the frequency of independent Cas3 translocation events, suggesting rupture between the Cse1 and Cas3 protein-protein contacts. (E) Top: illustration of the protein roadblock DNA substrate. Four EcoRI binding sites, E_1_ to E_4_, are positioned 4.8 kb, 10.4 kb, 17.9 and 23.7 kb upstream of the Cascade target. The hydrolytically defective EcoRI(E111Q) was used as a model protein roadblock. Bottom: kymographs showing outcomes of collisions between translocating Cascade/Cas3 complexes (magenta) and EcoRI(E111Q) (green). In all examples, collisions are shown with the first EcoRI(E111Q) bound at E1. (F) Quantification of the collision outcomes observed in (E). (G) Cascade/Cas3 translocation velocities (left) and processivities (right) on naked DNA and with EcoRI(E111Q) or LacI protein roadblocks. Red diamonds indicate the mean of the distribution. For experiments with LacI, the DNA substrate harbored a single ideal LacO site 12.3 kb upstream of the Cascade target. Dashed lines indicate the locations of E1 to E4 and the red lines indicate the location of the first roadblock encountered by Cascade/Cas3. N > 25 for all conditions. The translocation rate was statistically indistinguishable for all conditions (p=0.08, 0.34, 0.42 for EcoRI.E1, EcoRI.E2, and LacI relative to naked DNA, respectively), whereas the processivity was significantly reduced in all roadblock experiments (p=5.7×10;^−20^, 5.9×10^−19^, 1.6×10^−12^ for EcoRI.E1, EcoRI.E2, and LacI relative to naked DNA, respectively). (H) Model summarizing how Cascade/Cas3 translocates on crowded DNA. Cascade/Cas3 extrude a DNA loop while translocating processively until a collision with a protein roadblock (red octagon). Cascade/Cas3 either slip back or stall at the roadblock. Cas3 can also separate from Cascade. An independently translocating Cas3 can push and evict the roadblock from DNA.

In the cell, the ssDNA generated via Cas3 helicase and nuclease activities will be rapidly bound by single-stranded DNA binding protein (SSB). We therefore determined whether SSB regulates Cas3 activities, and used SSB-GFP to image the resulting ssDNA (Figure S6). The intensity of one SSB tetramer on a short ssDNA overhang was used to estimate the number of SSBs associated with each Cas3 (Figure S6A,B). Interestingly, SSB-GFP signal accumulated at Cascade/Cas3 complexes prior to processive translocation (Figure S6C). These puncta were not observed when either Cas3 or ATP were omitted from the flowcells (data not shown). The Cas3 and ATP-dependent generation of ssDNA suggests that Cas3 was translocating distances that were below the ~500 bp resolution of these assays. In agreement with this hypothesis, we occasionally observed repetitive >500 bp Cas3 translocation and slipping that was coincident with a growing SSB-GFP signal (Figure S6C). Consistent with our observations, a smFRET study also observed *Ec*Cas3 loop release and re-looping during translocation (Figure 3E), suggesting this is a universal feature of Type I-E Cas3 enzymes^57^. Moreover, the SSB-GFP signal only increased moderately during processive Cas3 translocation, and never reached full SSB saturation that would be expected if dsDNA were converted to ssDNA, suggesting Cas3 produces short tracts of ssDNA (N=36; Figure S6D). These results do not stem from SSB inhibition of Cas3, as neither velocity nor processivity were reduced with SSB added to the flowcell (Figure S6E). Taken together, our results are consistent with initiation via repetitive rounds of Cas3 slipping and restart, followed by processive Cas3 helicase activity and reannealing of the ssDNA into dsDNA. The Cas3 nuclease domain likely nicks the double-stranded DNA substrate and occasionally produces short tracts of ssDNA that are rapidly coated by SSB.

### Translocating Cascade/Cas3 is blocked by other DNA-binding proteins

In the cell, DNA is decorated with transcription factors and other DNA-binding proteins. Cas3 will likely encounter these obstacles during processive (>10 kb) translocation. We therefore determined if two site-specific DNA binding proteins—hydrolytically defective EcoRI(E111Q) and Lac repressor (LacI)—influence processive Cas3 translocation (Figure 3E-G). Both EcoRI(E111Q) and LacI bind their target sites with pM-nM affinity, and are frequently used as model roadblocks on DNA^59^. We first observed Cas3 interactions with fluorescent EcoRI(E111Q), which bound specifically to the four EcoRI binding sites on this DNA substrate. The closest two sites, EcoRI.E1 and EcoRI.E2, are 4.8 kb (E_1_) and 10.4 kb (E_2_) upstream of the Cascade target, respectively (Figure 3E, top). To assay Cas3 vs. EcoRI(E111Q) collisions, fluorescent Cascade and EcoRI(E111Q) were incubated with the DNA prior to assembling DNA curtains. Cas3 was introduced with ATP, and translocation was monitored via imaging of the Cascade/Cas3 looping complex. EcoRI(E111Q) blocked 100% (N=76/76 molecules) of all Cascade/Cas3 complexes. The most frequent outcome, accounting for 51% of all collisions (N=39/76), was Cascade/Cas3 stalling at the roadblock (Figures 3E,F). Other outcomes included stalling followed by dissociation of Cas3 from Cascade (33%), or single-frame release of Cascade/Cas3 back to the initial target site and re-looping by the same Cascade/Cas3 complex (8%). In the rare event of Cas3 dissociation from Cascade, the freely-moving Cas3 could push EcoRI(E111Q) off its target site (observed in 8% of Cascade/Cas3 collisions). We never observed roadblock pushing by the entire Cascade/Cas3 complex, suggesting that Cas3 alone may be more active at removing protein roadblocks. To differentiate the effects of the roadblock from the natural processivity of Cascade/Cas3 on naked DNA, we focused our analysis on Cascade/Cas3 complexes that encountered either of the first two occupied EcoRI(E111Q) binding sites. The observed velocity was statistically indistinguishable from Cas3 on naked DNA. However, translocation was blocked by the protein roadblock (Figure 3G, S7). LacI, located 12.3 kb upstream of the Cascade target, also acted as a strong roadblock to Cascade/Cas3 translocation, with 100% (N=28/28) of collisions resulting in stalling and frequent Cascade/Cas3 loop release (Figure S7). In sum, the Cascade/Cas3 complex processively translocates on naked DNA, but is blocked by other DNA-binding proteins (Figure 3H). Roadblocks may promote Cas3 slipping and re-looping, as has been observed in this study and with *Ec*Cas3^57^. Cascade/Cas3 may also stall frequently during translocation on crowded DNA *in vivo.* This stalling may provide additional time for the Cas3 nuclease activity to degrade foreign genetic elements and may also explain the degradation and primed acquisition hotspots reported in prior *in vivo* studies^5,11,39^. Stochastic rupture of the Cse1-Cas3 interface will eventually liberate freely-translocating Cas3 to push and evict roadblocks during CRISPR interference (Figure 3H).

### Cas1-Cas2 associates with Cascade/Cas3 in the Primed Acquisition Complex (PAC)

Primed acquisition requires Cascade, Cas3, and the Cas1-Cas2 integrase^5–12^. However, the functions of Cas1-Cas2 in primed acquisition have only been assayed indirectly. Here, we observed the assembly and translocation of a ~710 kDa primed acquisition complex (PAC), consisting of *Tfu* Cas1-Cas2, Cascade, and Cas3 (Figure 4). For single molecule imaging, the Cas2 N-terminus was fluorescently labeled via sortase-mediated transpeptidation (Figure S8). As expected from biochemical and structural studies of diverse Cas1-Cas2 integrases, *Tfu*Cas1-Cas2 also formed a heterodimer with a (Cas1)_4_-(Cas2)_2_ stoichiometry (Figure S8B). Cas1-Cas2 transiently bound the DNA substrate with a half-life of ~5.9 ± 0.1 seconds (N=38) (Figures 4A and 4D). We next sought to determine how Cas1-Cas2 interacts with the Cascade surveillance complex (Figure 4B). Three lines of evidence indicate that Cas1-Cas2 forms a long-lived complex with both target-bound and diffusing Cascade complexes. First, Cas1-Cas2 co-localized with Cascade that was pre-loaded on the target site and the lifetime of Cas1-Cas2 on DNA increased ~5.8-fold relative to Cas1-Cas2 in the absence of Cascade (Figures 4B and S8E). Second, pre-incubating fluorescent or unlabeled Cascade with fluorescent Cas1-Cas2, resulted in Cascade/Cas1-Cas2 complexes that diffused on non-specific DNA and could recognize the Cascade target sequence (Figure S8F). Third, Cascade could be pulled down with bead-immobilized *Tfu*Cas1-Cas2 (Figure S8G). Next, unlabeled Cas3 was added to the pre-assembled Cascade/Cas1-Cas2 sub-complex and the entire PAC was imaged via dual-color illumination. Directional translocation of the PAC away from the target site confirmed the presence of Cas3 (Figure 4C). Importantly, the PAC remained stationary when ATP was substituted for the non-hydrolyzable AMP-PNP in the imaging buffer (data not shown). The majority of translocating PACs retained Cas1-Cas2 for the duration of the entire trajectory (87.5%, N=35/40), indicating that Cas1-Cas2 is further stabilized within the PAC (Figures 4D & 4E) relative to the Cascade/Cas1-Cas2 sub-complex. All translocating PACs moved towards the DNA tether at a mean velocity of 84 ± 28 bp s^−1^ (N=40; error indicates S.D.), which was statistically indistinguishable from the velocity observed for Cascade/Cas3 (Figure 4F). In contrast, the PAC processivity was 20% lower than the Cascade/Cas3 complex (15.5 ± 5.6 kb for the PAC, N=40; p=0.015 relative to Cascade/Cas3). Whereas ~50% of Cascade/Cas3 complexes eventually showed Cse1-Cas3 rupture and independent Cas3 translocation, we did not see any independently translocating Cas1-Cas2/Cas3 sub-complexes under identical force and imaging conditions (Figure 4G, N=40). Taken together, these results suggest that the Cas1-Cas2 is a core subunit of PAC, where it is stabilized by direct interactions with Cascade. Additional interactions between Cas1-Cas2 and Cas3, as well as the forked DNA that emerges from the Cas3 exit channel may also contribute to Cas1-Cas2 retention in the PAC.

**Fig. 4.**
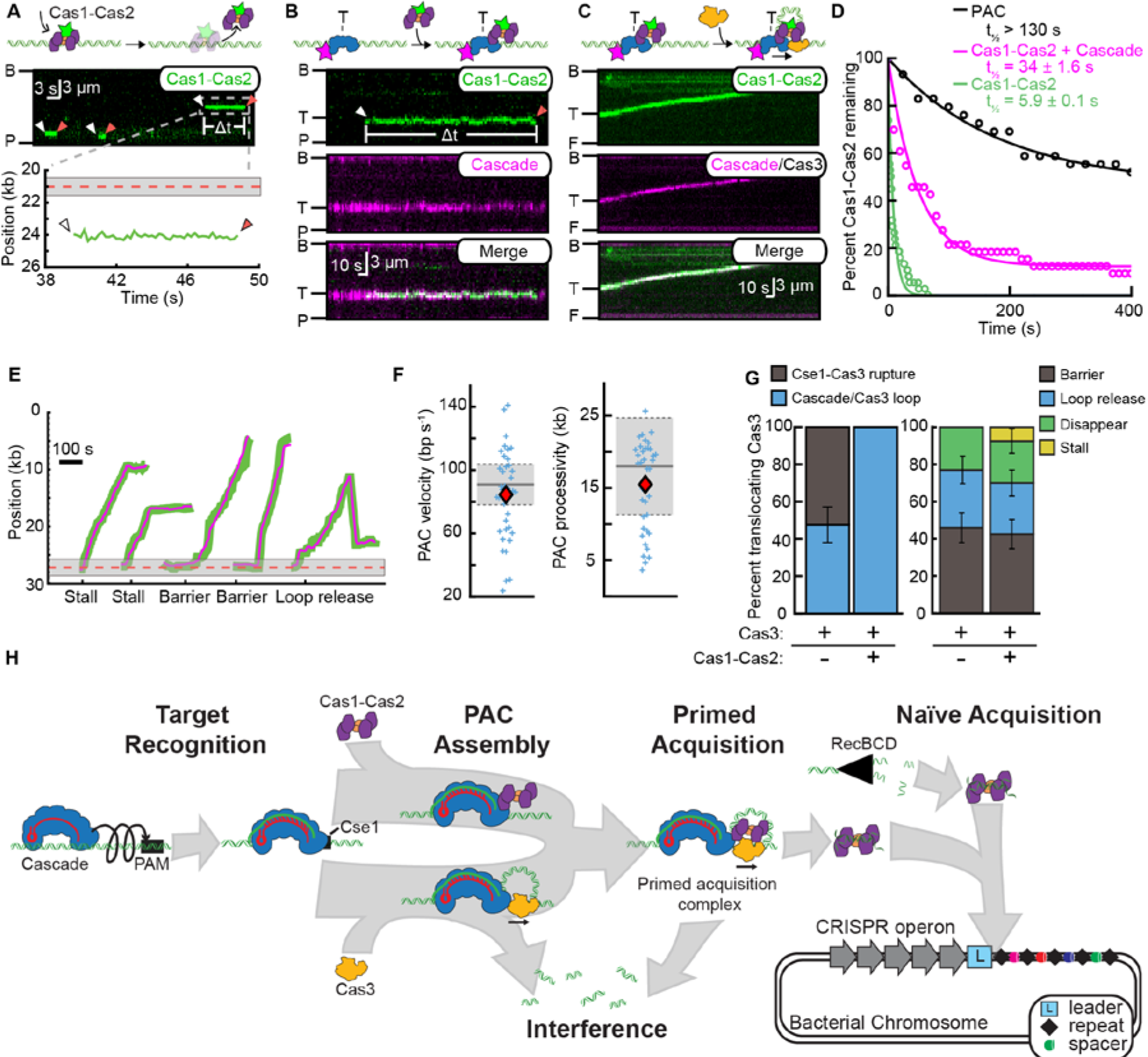
Cas1-Cas2 forms a primed acquisition complex (PAC) with Cascade and Cas3. (A) Illustration (top), kymograph (middle), and quantification (bottom) showing Cas1-Cas2 randomly sampling DNA via 3D collisions. White arrows: Cas1-Cas2 binding, red arrows: Cas1-Cas2 dissociation. The dashed red line and gray band represent the Cascade target site, as defined in Figure 2. (B) Illustration (top) and kymographs of Cas1-Cas2 (green) recruitment to Cascade (magenta) at the target sequence. (C) Illustration and a kymograph of the primed acquisition complex (PAC) consisting of Cascade, Cas1-Cas2, and Cas3 processively translocating along the DNA. Cascade (magenta) and Cas1-Cas2 (green) are fluorescently labeled while the presence of dark Cas3 is observed via translocation of the entire complex. (D) DNA-binding lifetimes of Cas1-Cas2 on DNA (green), with Cascade (magenta), and within the PAC (black). The data is fit to a single exponential decay. A constant was also included in the Cascade/Cas3 and PAC fits. Error: 95% C.I. (E) Representative traces of the PAC translocating on DNA. Cascade (magenta) and Cas1-Cas2 (green) are fluorescently labeled. (F) The mean PAC velocity (red diamond) was statistically indistinguishable from Cascade/Cas3 (N≥39; p=0.34). Mean PAC processivity was reduced compared to Cascade/Cas3 (p=0.015). Red diamonds indicate the mean of the PAC distribution. The mean and S.D. of the Cascade/Cas3 distributions are indicated by the solid and dashed lines, respectively. (G) Left: The PAC translocates exclusively via a DNA looping mechanism. Right: termination outcomes for translocating Cascade/Cas3 and PAC complexes. Error bars generated via bootstrapping. (H) Overview of the stepwise assembly of CRISPR-associated sub-complexes in interference and spacer acquisition. First, Cascade surveils foreign DNA via a combination of facilitated 1D diffusion and 3D collisions. Cse1 validates the PAM and initiates directional R-loop propagation, followed by additional conformational locks along the Cse2 subunits. Cascade can also recruit either Cas3 or Cas1-Cas2. The primed acquisition complex (PAC) consisting of Cascade, Cas1-Cas2, and Cas3 scans the foreign DNA for possible protospacers during processive translocation.

## Discussion

Here, we directly observe the first steps of target recognition and processing by the *Tfu* Type I-E CRISPR/Cas complex (**Figure 4H**). Cascade scans non-specific DNA for its target via a combination of 1D facilitated diffusion and 3D collisions. An evolutionarily-conserved positive patch on the outer surface of Cse1 promotes facilitated diffusion of Cascade during target recognition. Neutralizing mutations in this positive patch reduce the lifetimes of diffusing Cascade complexes on non-specific DNA and decrease the binding affinity for the target DNA 50-fold. Target recognition and stable R-loop locking proceeds via at least two temporally distinct intermediates. The first of these intermediates initiates PAM-proximal opening of the DNA bubble and sampling of the target DNA “seed” region. The second, longer-lived intermediate includes R-loop propagation and additional stabilization via Cse2-dependent salt-bridges. Abortive complexes that cannot fully recognize the R-loop dissociate from the DNA target and continue to scan nonspecific DNA. After target recognition, Cascade recruits Cas3 helicase/nuclease and the Cascade/Cas3 complex translocates in a 3’→5’ direction on the non-target strand. Cascade remains associated with the target DNA, causing a growing nicked DNA loop to develop between the Cas3 motor and a target-bound Cascade. This protein interaction ruptures in a stochastic and force-dependent manner, with Cas3 continuing to translocate independently of Cascade. The Cascade/Cas3 complex is highly processive on naked DNA but is blocked by other DNA-binding proteins. In contrast, after Cas3 separates from Cascade, the freely-moving motor can push protein roadblocks from their DNA-binding sites. Clearing protein roadblocks by Cas3 can improve the interference efficiency and may allow uptake of new protospacers further away from the initial target sequence. Primed acquisition also requires the Cas1-Cas2 integrase. Here, we provide the first direct evidence that Cas1-Cas2 is stabilized on DNA via physical interactions with Cascade. Cascade forms the keystone of the PAC, as Cas3 and Cas1-Cas2 both require Cascade for stable association with the target DNA. Our data suggest that the PAC can assemble via two routes that include initial recruitment of either Cas3 or Cas1-Cas2 to target-bound Cascade, followed by addition of the remaining sub-complex (**Figure 4H**). Further support for this assembly comes from genetic, biochemical, and structural studies of the Type I-F system, where Cas3 is expressed as a direct fusion with Cas2^10,60^. Finally, we demonstrate that Cas1-Cas2 translocates with Cascade and Cas3 as part of the PAC. Cas1-Cas2 harbors a PAM-decoding center, initially identified in the structure of the *Ec*Cas1-Cas2 complex, and is also conserved in *Tfu*Cas1-Cas2 (**Figure S8H**)^35^. As part of the PAC, the Cas1-Cas2 PAM decoding center can scan, capture, and excise foreign DNAs as they emerge from Cas3. This likely uses the Cas1 nuclease active site, as the Cas2 nuclease is structurally occluded and dispensable for integration *in* vivo^35,61,62^. Moreover, the combined endonuclease activities of Cas3 and Cas1 may further stimulate efficient DNA degradation during PAC translocation and primed acquisition. Additional studies will be required to address precisely how Cas1-Cas2 selects protospacers during PAC translocation and how these protospacers are subsequently integrated into the bacterial genome.

## Materials and Methods

### Protein cloning and purification

*Thermobifida fusca (Tfu)* Cascade^56^, *Tfu*Cas3^56^, *E. coli (Eco)* SSB^63^, EcoSSB-GFP^64^, *Eco* 3xHA-EcoRI(E111Q)^64^, *Eco* 3xHA-LacI^64^, sortase variants^50,65,66^ and SUMO protease^67^ were purified as described previously. *Tfu*Cas3 was also purified using a M9 minimal media excluding trace metals. *Tfu* Cas1 and Cas2 were cloned into pET expression vectors containing an N-terminal His_6_-SUMO-fusion. Cas1 and Cas2 were purified separately, reconstituted as a complex, and gel filtered over a HiPrep Sephacryl S-200 HR column.

### Sortase labeling for single-molecule imaging

Peptides were synthesized using the Liberty Blue Automated Microwave Peptide Synthesizer (CEM Corporation) using manufacturer-suggested protocols, and labeled with NHS-Atto647N (Atto-Tec). For fluorescent labeling, Cse1 and Cas2 were purified with an N-terminal GGG residues after the SUMO tag and Cas3 was purified with a C-terminal LPETGG-TwinStrep motif. Sortase labeling was optimized for each protein by varying the temperature, labeling time, and sortase variant^50,65,66^. Labled proteins where isolated using a HiPrep Sephacryl S-200 HR column (GE) with TS Buffer (10 mM Tris-HCl [pH 7.5], 150 mM NaCl, 5 mM DTT).

### DNA substrates for single-molecule microscopy

DNA substrates with mutated target sequences were generated by cloning the mutated targets into helper plasmids pIF152 and pIF153 that had ~200 bp of flanking homology with λ-phage DNA^68^. To functionalize the DNA ends for single-molecule experiments, we combine 125 μg of purified λ-phage DNA with 2 μM of biotinylated oligos. For double-tethered DNA curtains, a second dig-labeled oligo was annealed to the second DNA end. After ligation, the reaction was separated over a Sephacryl S-1000 column (GE, #45-000-084) to purify full length labeled DNA. The DNA was stored at 4°C.

### Single-molecule fluorescence microscopy

All single-molecule imaging was performed using a Nikon Ti-E microscope in a prism-TIRF configuration equipped with a motorized stage (Prior ProScan II H117) containing microfluidic flowells housed in a custom stage adapter. The flowcell was illuminated with 488 nm (Coherent), 532 nm (Ultralasers), and 633 nm (Ultralasers) lasers through a quartz prism (Tower Optical Co.)^41^. A custom built microscope stage heater were used to maintain the flowcell near the optimal *Tfu*Cascade temperature.

### Data analysis

Fluorescent particles were tracking using an in-house ImageJ script (available upon request). Trajectories were used to calculate the mean-squared displacement and the diffusion coefficients for Cascade, or the velocity and processivity for the Cas3-containing complexes, as described previously ^69,63^. Binding lifetimes were fit to either a single exponential decay or a biexpoential decay using a custom MATLAB script (Mathworks R2015b). The biexpoential fits were tested to be appropriate using an *f-test* applied to the survival curve data^63^. For pause analysis, a molecule was considered paused if it stayed within a stationary window for four continuous frames (0.8 seconds). This window was defined as 3-fold the standard deviation (S.D.) of the fluctuations of a stationary Cascade at its target^47^. Pause location was recorded in relation to the pedestal located at the digoxigenin labeled end of the DNA.

Translocating Cas3 was defined as Cas3 that left the target window for at least four continuous frames (> 800 ms). Looping Cas3-Cascade molecules were defined by scoring whether Cascade also left the target window with Cas3. In contrast, independently moving Cas3s were defined by scoring traces where Cascade remained stationary while Cas3 moved away from the target window.

## Author Contributions and Notes

M.W.B., K.E.D., L.R.M., Y.X., A.K., and I.J.F. conceived the study. M.W.B., K.E.D., L.R.M., Y.X., A.K., and I.J.F. designed the experiments and analyzed the data. M.W.B., K.E.D., Y.X., A.D., and L.R.M. generated key materials and executed the experiments. E.H. and S.D. provided fluorescent peptides. Y.K. provided particle tracking and roadblock bypass software. M.W.B., K.E.D., and I.J.F. wrote the paper with input from all other authors.

The authors declare no conflict of interest.

## Acknowledgments

We are very grateful to Jeffrey Schaub, James R. Rybarski, Fatema A. Saifuddin, Andrew Leal, and Brianna Gonzalez for providing materials. This work was supported by the Welch Foundation (F-1808 to I.J.F.), the US National Institutes of Health grant R01GM124141 to I.J.F. and R35GM118174 to A.K. I.J.F. is a CPRIT Scholar in Cancer Research. Yoori Kim is a HHMI graduate student fellow. L.M. was supported by NIH fellowship F99CA212452.

